# Periplasmic coiled coil formed by assembly platform proteins PulL and PulM is critical for function of the *Klebsiella* type II secretion system

**DOI:** 10.1101/2022.09.21.508927

**Authors:** Yuanyuan Li, Javier Santos-Moreno, Olivera Francetic

## Abstract

Bacteria use type II secretion systems (T2SS) to secrete to their surface folded proteins that confer diverse functions, from nutrient acquisition to virulence. In the *Klebsiella* species, T2SS-mediated secretion of pullulanase (PulA) requires assembly of a dynamic filament called pseudopilus. The inner membrane assembly platform (AP) complex is essential for PulA secretion and pseudopilus assembly. The AP components PulL and PulM form an inner membrane complex interacting through their C-terminal globular domains and transmembrane segments. Here we investigated the roles of periplasmic helices and cytoplasmic domains of PulL and PulM in their assembly. We found that PulL and PulM variants lacking periplasmic helices were defective for interactions in the bacterial two-hybrid (BACTH) assay. Their function in PulA secretion and assembly of PulG subunits into pseudopilus filaments were strongly reduced. In addition, deleting the cytoplasmic peptide of PulM in variant PulMΔN nearly abolished interaction with PulG in the BACTH assay, without affecting the interaction with PulL. Nevertheless, PulL was degraded in the presence of the PulMΔN variant, suggesting that PulM N-terminal peptide interacts with PulL in the cytoplasm and plays a stabilizing role. We discuss the implication of these results for the mechanism of T2S and type IV pilus assembly.

## 1. Introduction

Bacteria have an astonishing capacity to adapt to diverse environments, growth conditions and nutrient sources. Understanding the molecular basis of this adaptation is fundamentally important and might lead to applications in biotechnology and bioremediation. Among the numerous nanomachines that contribute to bacterial adaptation, type II secretion system (T2SS) plays a prominent role (Cianciotto & White, 2017).

Recent reviews summarize our current knowledge of T2SS architecture and molecular function (Gu, Shevchik et al., 2017; Korotkov & Sandkvist, 2019; Naskar, Hohl et al., 2021). T2SS is typically composed of 15 proteins forming a megadalton complex that spans the envelope of Gram-negative bacteria. In the outer membrane, 15 subunits of the GspD protein form the secretin channel that allows the passage of large folded proteins or their complexes to the cell surface. According to the current models, protein secretion is driven by the polymerization of periplasmic filaments called pseudopili in the inner membrane. The core of the pseudopilus is a helical homopolymer composed of the major subunit GspG (Sauvonnet, Gounon et al., 2000; Durand, Bernadac et al., 2003; Kohler, Schafer et al., 2004). The minor pseudopilins GspH, I, J and K from a complex at the pseudopilus tip (Korotkov & Hol, 2008; Douzi, Ball et al., 2011) and initiate pseudopilus assembly (Cisneros, Bond et al., 2012). The minor pseudopilins are essential for protein secretion and have also been implicated in substrate recognition (Douzi et al., 2011). Pseudopili are continuously assembled by the inner membrane at the assembly platform (AP) subcomplex composed of GspF, GspL and GspM proteins (Py, Loiseau et al., 2001). GspL forms a stable complex with the cytoplasmic ATPase GspE (Abendroth, Murphy et al., 2005) and is itself stabilized by GspM (Sandkvist, Hough et al., 1999; Possot, Vignon et al., 2000; Lallemand, Login et al., 2013). During pseudopilus elongation, GspM promotes targeting of pseudopilins GspG and GspH to the AP (Nivaskumar, Santos-Moreno et al., 2016; Santos-Moreno, East et al., 2017). In the cytoplasm, cycles of ATP binding and hydrolysis by the GspE ATPase promote conformational changes of the GspE hexameric complex (Robien, Krumm et al., 2003; Patrick, Korotkov et al., 2011). These changes are proposed to induce rotational movements of GspF at the center of the AP that promote pseudopilus assembly (Nivaskumar, Bouvier et al., 2014).

Deciphering the structural basis of pseudopilus assembly is important to understand how T2SS functions at the molecular level. Structure-function analyses have provided molecular insight into the majority of individual T2SS components. Extending this knowledge towards larger subcomplexes is bringing clues on the connectivity between these components. Cryo-electron microscopy has been essential to gain insight into the secretin channel structure (Yan, Yin et al., 2017; Hay, Belousoff et al., 2018; Chernyatina & Low, 2019), a large pentadecameric complex with a periplasmic gate. Combining cryoEM, NMR, modeling and mass spectrometry provided a detailed view of the pseudopilus core (Campos, Nilges et al., 2010; (López-Castilla, Thomassin et al., 2017) and the tip complexes (Escobar, Douzi et al., 2021).

Recent studies of the *Klebsiella oxytoca* T2SS that secretes pullulanase (PulA) combined X-ray crystallography and NMR to gain information on the structure of a complex formed between C-terminal domains (CTDs) of GspL (PulL) and GspM (PulM) (Dazzoni, Li et al., 2022). Mutations at the interface of these CTDs affect the stability of the two proteins and their function in protein secretion. The model of the PulL-PulM complex (lacking the cytoplasmic domain of PulL) predicts additional interactions *via* periplasmic helices of the two partners. Here we investigated the importance of this interface for the formation and function of the PulL-PulM complex. In addition, we studied the role of the short cytoplasmic region of PulM in the stability and function of the AP complex.

## 2. Materials and methods

### 2.1. Bacterial strains and culture conditions

*Escherichia coli* strain DH5αF’*lacI*^Q^ was used for cloning and transformation experiments. Functional assays were performed in *E. coli* PAP7460 (Possot et al., 2000) and PAP5378 strains. PAP5378 was constructed by P1 transduction by introducing the *ompT::kan* gene from the Keio collection strain BW2511 *ompT::kan* into strain PAP5299 (Cisneros et al., 2012). The *kan* cassette was then deleted thanks to the helper plasmid pCP20 as described (Datsenko & Wanner, 2000) to give strain PAP5299 *DompT*. The bacterial two-hybrid assays were performed in strain DHT1 (Dautin, Karimova et al., 2000). Bacteria were grown at 30°C or 37°C in LB medium (Miller, 1972) supplemented with antibiotics as required: ampicillin (Ap) at 100 μg.mL^-1^, chloramphenicol (Cm) at 25 μg.mL^-1^, Kanamycin (Km) at 25 μg.mL^-1^. Solid LB media contained 1.5% agar (Difco). Expression of genes under *lacZ* promoter control was induced with 1 mM isopropyl-β-1-D-thiogalactopyranoside (IPTG). The *pul* gene promoters were induced with 0.4% D-maltose. For secretion assays, media were buffered with 0.1 volume of M63 salt solution (Miller, 1972).

### 2.2. Plasmid construction and site-directed mutagenesis

Plasmids used in this study are listed in Table 1. Site-directed mutagenesis was performed using a modified QuickChange protocol, with fully overlapping oligonucleotide primers listed in Table S1 (Eurofins). Template DNA was amplified with high fidelity DNA polymerase Q5 (New England Biolabs) in two-steps: first, 6 cycles of amplification were performed with forward and reverse primers alone. The reactions were mixed and another 15 cycles of amplification were performed. The DNA was treated with *DpnI* enzyme (Fermentas) and transformed into DH5αF’ ultracompetent cells. Isolated colonies were inoculated into 5 mL of LB with appropriate antibiotics and plasmids were purified using the alkaline lysis protocol and Qiaquick miniprep kit (Qiagen). Plasmids were verified by sequencing (GATC, Eurofins). For the *pulL* bacterial two hybrid constructs, vectors pUT18C and pKT25 were digested with *Eco*RI and *KpnI* and the purified vector fragments were ligated with the PCR amplified *pulL Δ270-302* digested the same enzymes. For the *pulM* cloning, the BACTH vectors and PCR fragments were digested with *Eco*RI and *Bam*HI. Deletion of *pulM* residues 2-15 in plasmid pCHAP8882 was performed by QuickChange mutagenesis using primers PulM delN5 and PulM delN3 (Table S1). The same approach was used to introduce PulM2-15 deletion in the BACTH constructs pCHAP8154 and pCHAP8155 with primers PulM delN For and PulM delN Rev2 (Table S1).

**Table 1.** Plasmids used in this study.

| **Name** | **Relevant characteristics** | **Source/Reference** |
| --- | --- | --- |
| pCHAP8185 | *pulC-O, pulAB, pulS, Ap<sup>R</sup>, ColE1ori* | (Cisneros et al., 2012) |
| pCHAP8496 | pCHAP8185 Δ*pulM* | (Santos-Moreno et al., 2017) |
| pCHAP8251 | pCHAP8185 Δ*pulL* | (Santos-Moreno et al., 2017) |
| pCHAP8258 | *placZ-pulL, Cm<sup>R</sup>, p15A ori* | (Santos-Moreno et al., 2017) |
| pMS1349 | pCHAP8258 *pulL* Δ270-302, Cm<sup>R</sup>, p15A *ori* | This study |
| pCHAP1353 | *placZ-pulM, Cm<sup>R</sup>, p15A ori* | (Possot et al., 2000) |
| pMS1350 | pCHAP8258 *pulM*Δ37-68, Cm<sup>R</sup>, p15A *ori* | This study |
| pCHAP8882 | *pulM*Δ2-15 (*pulM*ΔN) Cm<sup>R</sup>, p15A *ori* | This study |
| pUT18C | *cyaT18 Ap<sup>R</sup>, ColE1ori* | (Karimova et al., 1998) |
| pKT25 | *cyaT25 Km<sup>R</sup>, p15A ori* | (Karimova et al., 1998) |
| pMS1222 | *T18-pulL Ap<sup>R</sup>, ColE1ori* | (Dazzoni et al., 2022) |
| pMS1229 | *T25-pulL, Km<sup>R</sup>, p15A ori* | (Dazzoni et al., 2022) |
| pCHAP8154 | *T18-pulM, Ap<sup>R</sup>, ColE1ori* | (Nivaskumar et al., 2016) |
| pCHAP8155 | *T25-pulM, Km<sup>R</sup>, p15A ori* | (Nivaskumar et al., 2016) |
| pMS1355 | *T18-pulL*Δ270-302, Ap<sup>R</sup>, ColE1*ori* | This study |
| pMS1356 | *T25-pulL*Δ270-302, Km<sup>R</sup>, p15A *ori* | This study |
| pMS1357 | *T18-pulM*Δ37-68 Ap<sup>R</sup>, ColE1*ori* | This study |
| pMS1358 | *T25-pulM*Δ37-68 Km<sup>R</sup>, p15A *ori* | This study |
| pCHAP7330 | *T18-pulG, Ap<sup>R</sup>, ColE1ori* | (Nivaskumar et al., 2014) |
| pCHAP7332 | *T25-pulG, Km<sup>R</sup>, p15A ori* | (Nivaskumar et al., 2014) |
| pMS1147 | *T18-pulM*Δ2-15 Ap<sup>R</sup>, ColE1*ori* | This study |
| pMS1148 | *T25-pulM*Δ2-15 Km<sup>R</sup>, p15A *ori* | This study |

### 2.3. Protein electrophoresis and Western blot analysis

Total bacterial extracts were analyzed on denaturing sodium dodecyl-sulphate polyacrylamide gel electrophoresis (SDS-PAGE) using the Tris-tricine gels (Schägger & von Jagow, 1987). Proteins were transferred on nitrocellulose membranes using the Fast Blot semi-dry transfer system and one-step transfer buffer (Thermo). Membranes were blocked with 5% skim milk in Tris-buffer saline containing 0.05% Tween-20 (TBST) for one hour and probed with primary antibodies diluted in TBST. Anti-PulM and anti-PulL sera were used at 1:1000 dilution and anti-PulA and anti PulG sera were diluted at 1:2000. After 4 washes for 10 minutes in TBST, membranes were incubated 1 hour in the secondary goat anti-rabbit antibodies coupled to horse-radish peroxidase (Amersham) (1:10000). Fluorescence signals were developed with ECL2 (Thermo) and recorded on Typhoon FLA9000 (GE, Cytiva).

### 2.4. PulA secretion assays

Secretion of the non-acylated variant of PulA was assessed by cell fractionation in strain PAP7460 harboring plasmid pCHAP8251 complemented with pCHAP8258 derivatives carrying *pulL* variants or pCHAP8496 complemented with pCHAP1353 derivatives carrying *pulM* variants. Bacteria were cultured overnight at 30°C in LB containing Ap and Cm, then inoculated in inducing medium containing in addition 0.4 % D-maltose and 0.1 vol of M63 salts for another 4 hours. Cultures were normalized to OD_600nm_ of 1. Bacterial pellets were collected after 5-min centrifugation at 16000 x g in a table-top Eppendorf centrifuge at 4°C and resuspended in an equal volume of Laemmli sample buffer. Supernatants were subjected to another 10-min round of centrifugation and 0.1 ml was taken off top, then mixed with 0.1 ml of 2 x Laemmli sample buffer. Cell and supernatant fractions from the same amount of cultures were analyzed by SDS-PAGE on 10% Tris-glycine gels followed by Western blot with anti-PulA antibodies. PulA bands were quantified with ImageJ and the fraction of PulA in the supernatant was calculated. Data were plotted and analyzed with GraphPad Prism9 software.

### 2.5. PulG pilus assembly assays

To quantify the PulG pilus assembly, bacteria of strain PAP7460 harboring the *pul* operon on plasmid pCHAP8185 derivatives were cultured for 48 or 72 hours at 30°C on LB plates containing 1.5% agar (Difco), appropriate antibiotics and 0.2% D-maltose. Bacteria were resuspended in 1 ml of LB and the cell density was normalized to OD_600nm_ of 1. The suspensions were vigorously vortexed for 1 min to detach surface pili. Bacterial pellets were then collected by 5-min centrifugation at 4°C and 16000xg and resuspended in Laemmli sample buffer at the concentration of 10 OD_600nm_.mL^-1^. The pili-containing supernatants were cleared from remaining bacteria by a 10-min centrifugation at 16000xg and 0.7 mL was mixed with tri-chloro-acetic acid (TCA) at a final concentration of 10%. After a 30-min precipitation on ice, the samples were centrifuged for 30 min at 4°C and 16000xg. Pellets were washed twice with acetone and air dried, then resuspended in 70 μL of Laemmli sample buffer. Equivalent volumes were analyzed by SDS-PAGE on 10 % Tris-tricine gels followed by Western blot with anti-PulG antibodies. PulG bands were quantified using ImageJ and the results were analyzed and plotted with Prism 9 software (GraphPad).

### 2.6. Bacterial two-hybrid assays

Bacterial two-hybrid (BACTH) assay was used to study interactions between proteins fused to T18 and T25 fragments of the CyaA catalytic domain (Karimova, Pidoux et al., 1998). Plasmids were co-transformed into DHT1 competent cells and grown on LB Ap Km plates for 48-60 hours at 30°C. Independent single co-transformants were picked at random, inoculated into 1 mL of LB Ap Km and grown overnight at 30°C. The overnight precultures were used to inoculate 1 mL cultures of LB Ap Km containing 1 mM IPTG, which were incubated 4-6 hours at 30°C with shaking. The cultures were used to measure beta-galactosidase activity as described by (Miller, 1972). The data were plotted and analyzed using the Prism 9 software (GraphPad).

## 3. Results

### 3.1. The periplasmic helices of PulL and PulM are important for T2SS function

PulL and PulM share a similar architecture of their transmembrane and periplasmic domains (Fig. 1A). Their hydrophobic transmembrane helices extend into the periplasm and are followed by flexible linkers connected to the globular ferredoxin-like C-terminal domains (CTDs). The two periplasmic helices are predicted to form a coiled coil (Dazzoni et al., 2022). To test their functional importance, we generated PulL and PulM variants lacking the periplasmic helices. In PulL, we removed the region comprising residues 270 to 302 (Fig. 1A) to yield variant PulL^ΔCC^, encoded by plasmid pMS1349 (Table 1). We first tested the ability of PulL^ΔCC^ to promote secretion of the nonacylated variant of pullulanase (PulA), the substrate of the *K. oxytoca* T2SS (Michaelis, Chapon et al., 1985). Whereas the native PulL (PulL^WT^) promoted efficient PulA secretion, the majority of the PulA pool accumulated in the cell fraction in the presence of PulL^ΔCC^ (Fig. 1B). The accumulation of intracellular PulA was complete in the negative control lacking PulL. We used antibodies directed against PulL^CTD^ to assess PulL stability in these strains (Fig. 1B, lower panel labeled CCF). Quantification of PulL^ΔCC^ signals showed that protein levels were reduced by about 30% compared to the native PulL (Fig. 1D).

**Figure 1.**
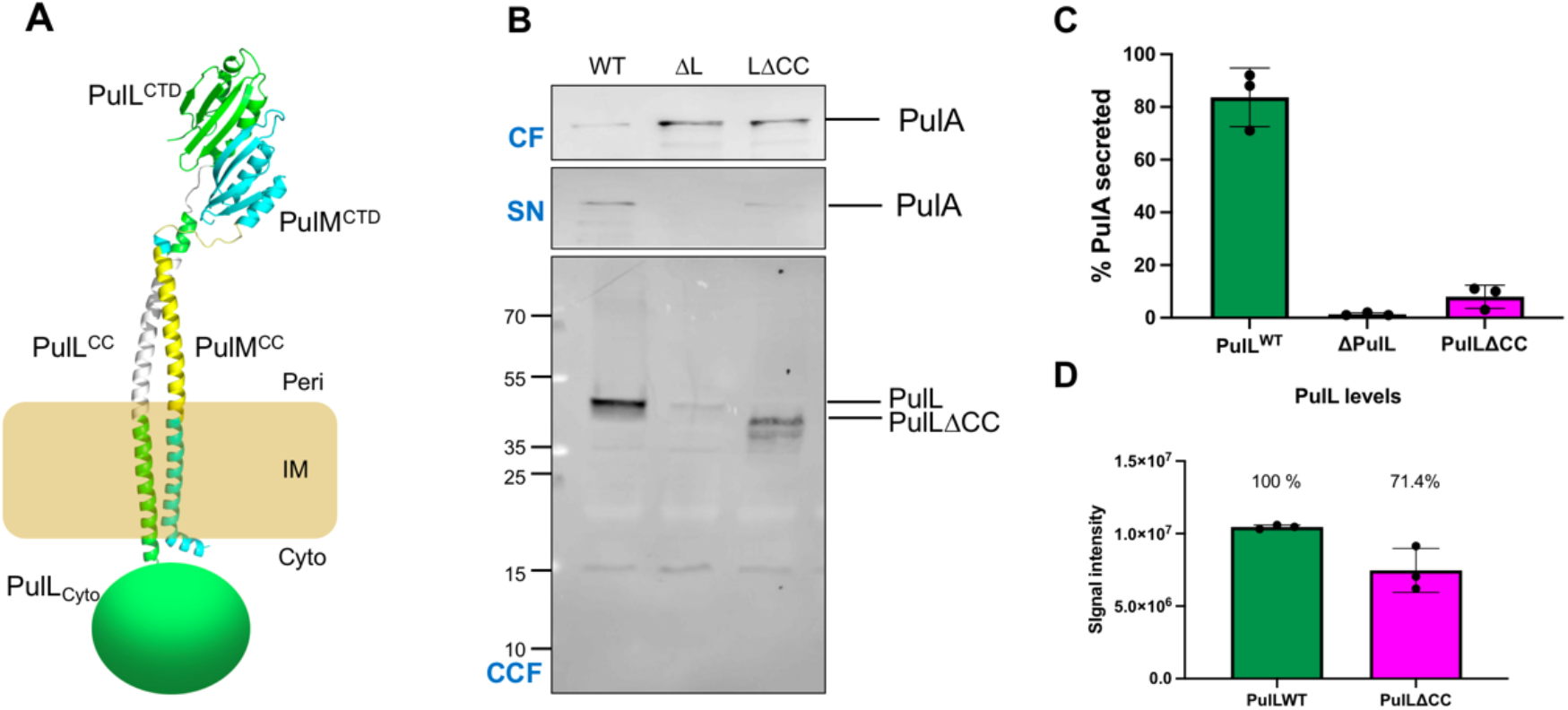
Deletion of the PulL periplasmic helix affects PulA secretion. **A**. Schematic view of the PulL-PulM dimer in the inner membrane (IM). PulL is shown in green with its periplasmic helix in white; PulM is shown in cyan and its periplasmic helix in yellow. **B**. Secretion of PulA in strains producing native PulL (WT), no PulL (ΔL) and PulL^ΔCC^. PulA secretion assay was performed (Materials and Methods) and 0.005 OD_600nm_ equivalents of cell- (CF) and supernatant fractions (SN) were analyzed by Western blot with anti-PulA antibodies. In the lower panel, the concentrated cell fractions (CCF) (from 0.05 OD_600nm_ of bacteria) were analyzed with anti-PulL antibodies. **C**. PulA secretion efficiency quantified from 3 independent assays as in (**B**). The column height shows the mean values and dots represent values from independent experiments. **D**. Bar graphs indicate mean signal intensities of bands detected with anti-PulL antibodies in strains producing PulL^WT^ and PulL^ΔCC^. Mean levels of PulL^ΔCC^ signal relative to PulL^WT^ are indicated above the bar. Black dots indicate signal levels from 3 independent measurements and error bars show standard deviation.

To study the role of the periplasmic helix of PulM, we removed the region comprising residues 37 to 68 (Fig. 1A) resulting in variant PulM^ΔCC^ encoded by plasmid pMS1350 (Table 1). Similar to the PulL case, this deletion resulted in a strong PulA secretion defect (Fig. 2A, 2B). PulA accumulated in the cell fraction in the presence PulM^ΔCC^ at levels similar to the negative control (Fig. 2A). PulM ^ΔCC^ stability was reduced by 10% on average compared to PulM (Fig. 2C).

**Figure 2.**
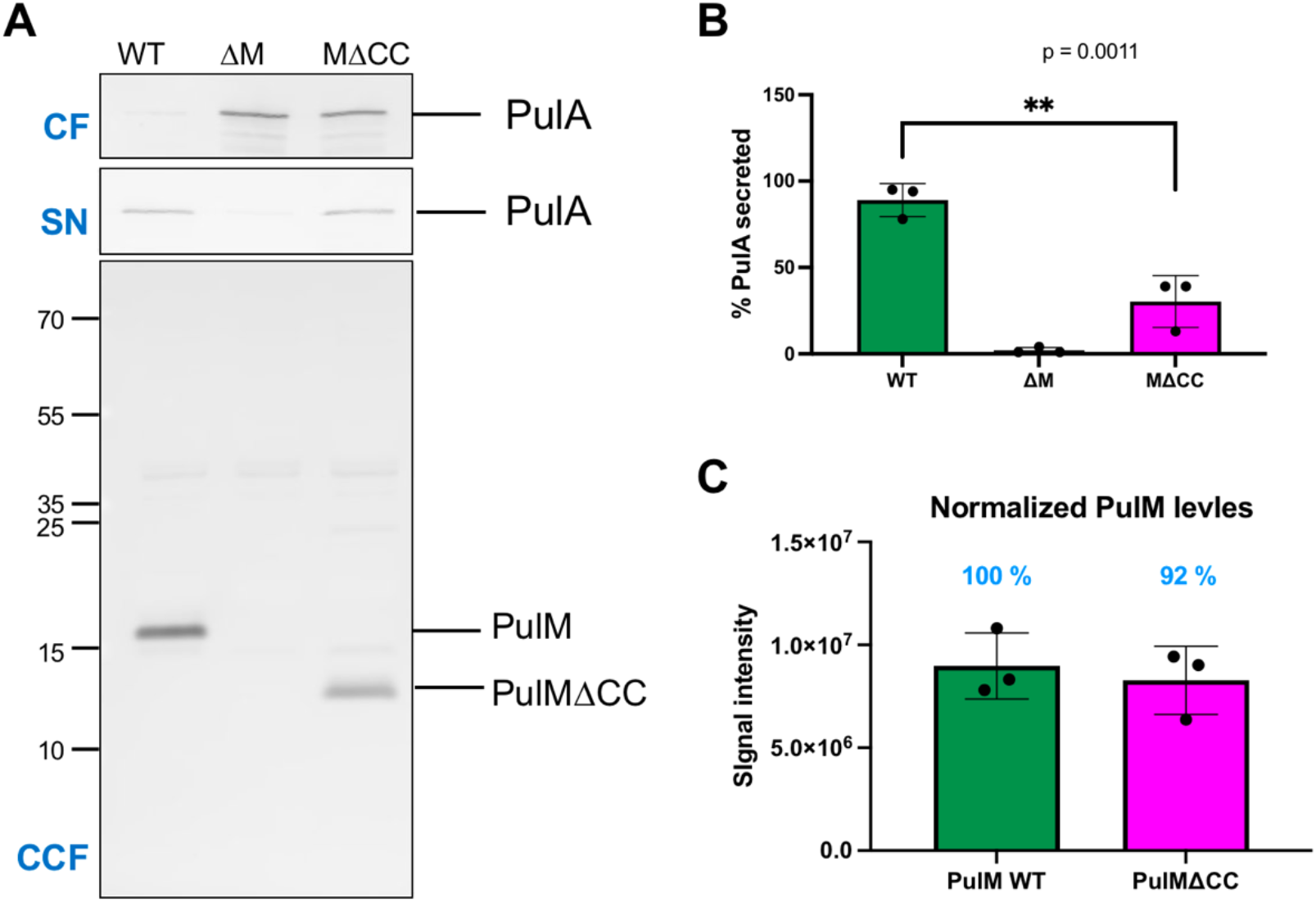
PulM periplasmic helix region is required for full T2SS function. **A**. Secretion of PulA in strains producing native PulM (WT), no PulM (ΔM) and PulM^ΔCC^. PulA secretion assay was performed (Materials and Methods) and 0.005 OD_600nm_ equivalents of cell- (CF) and supernatant fractions (SN) were analyzed by Western blot with anti-PulA antibodies. Lower panel, the concentrated cell fractions (CCF) (0.05 OD_600nm_) analyzed with anti-PulM antibodies. **B**. PulA secretion efficiency quantified from 3 independent assays in the presence of PulM (WT) or PulM^ΔCC^ and without PulM (ΔM). The column height shows the mean value and dots represent values from independent experiments. **C**. Bar heights indicate mean signal intensity of bands detected with anti-PulM antibodies in strains producing PulM^WT^ and PulM^ΔCC^. The mean fractions of relative signal intensity of PulM^ΔCC^ and PulM^WT^ are indicated above the bars. Black dots show signal intensities from 3 independent experiments and error bars show standard deviations.

### 3.2. *Pilus assembly defects in* pulL^ΔCC^ *and* pulM^ΔCC^ *mutants*

The AP components PulL and PulM directly participate in pseudopilus assembly, which is thought to drive PulA secretion. We therefore asked whether the periplasmic helices of PulL and PulM are required for PulG pilus assembly. Under conditions where bacteria are cultured on solid media, *E. coli* harboring a moderate copy-number plasmid encoding a complete set of *pul* genes produces surface pili composed of PulG, the major pseudopilin subunit (Sauvonnet, Vignon et al., 2000). We tested the ability of PulL^ΔCC^ and PulM^ΔCC^ to complement this function in mutants lacking plasmid-encoded *pulL* (pCHAP8251) or *pulM* (pCHAP8496). After 2 days of growth in the presence of maltose to induce the *pul* gene expression, bacteria complemented with wild type *pulL* and *pulM* genes produced pili that could be sheared from their surface and separated from the cell-associated PulG pool (Fig. 3). PulL^ΔCC^ and PulM^ΔCC^ variants allowed for weak secretion of PulA and the PulG pilus assembly was nearly abolished. When bacteria were cultured for a standard period of 48 hrs for this assay, no PulG could be detected in the pilus fractions.

**Figure 3.**
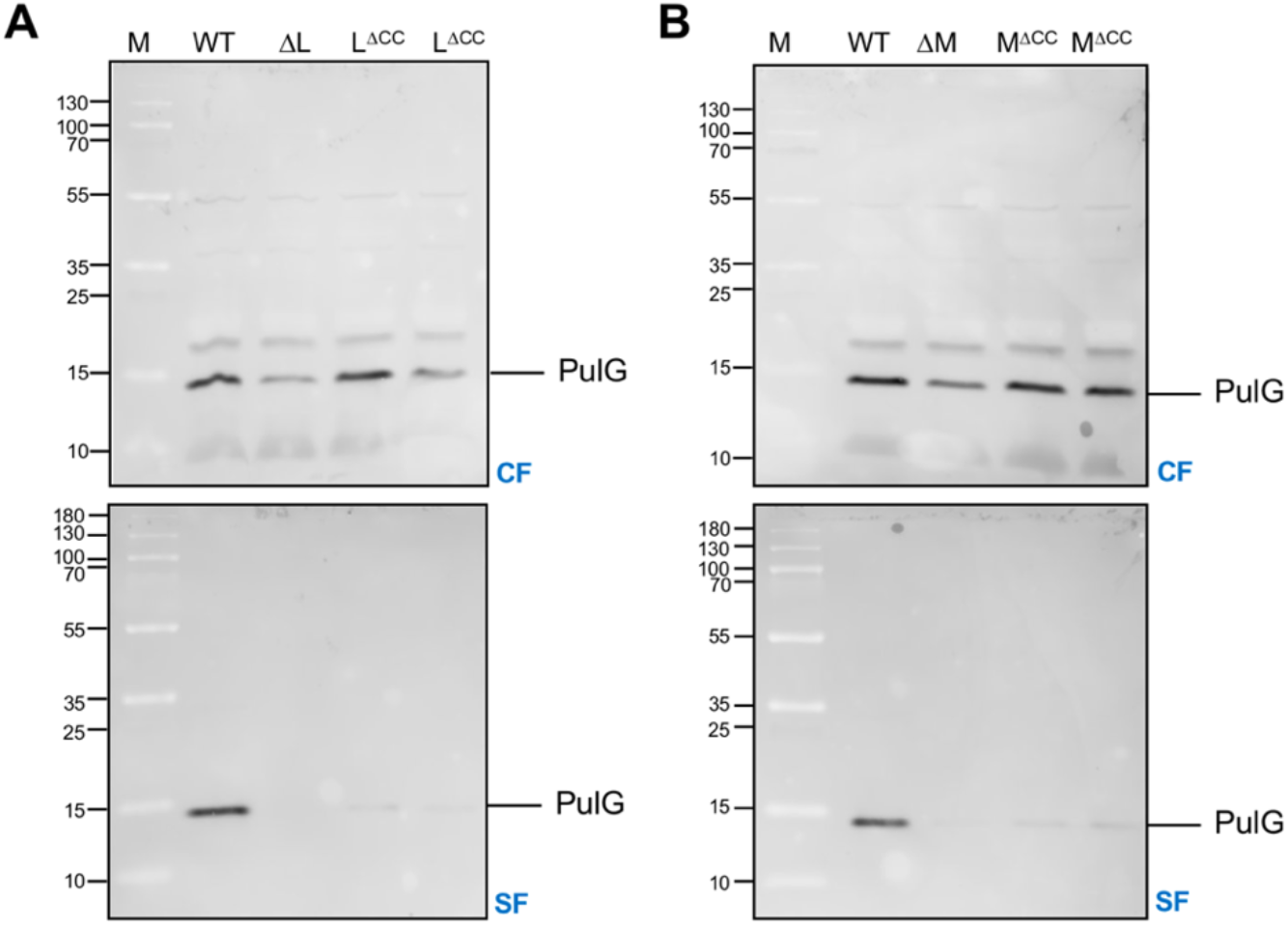
Piliation defect of PulL and PulM lacking periplasmic helices. **A**. PulG pilus assembly on the surface of PAP7460 bacteria containing plasmid pCHAP8251 complemented with pCHAP8252 (PulL^WT^), pMS1348 (PulL^ΔCC^) or vector pSU19 (ΔL). Migration of Mw markers (M) (in kDa) is indicated on the left. **B**. PulG pilus assembly on the surface of PAP7460 bacteria containing plasmid pCHAP8496 complemented with pCHAP1353 (PulM^WT^), pMS1350 (PulM^ΔCC^) or vector pSU18 (ΔM). Cell- (CF) and sheared fractions (SF) were prepared as described in Materials and Methods and 0.05 OD_600nm_ equivalents was analyzed on 10% Tris-Tricine SDS PAGE and western blot using anti-PulG antibodies.

Only upon extending the growth period for another 24 hours, we could detect some PulG pili on the surface of the coiled coil deletion mutants (Fig. 3). In addition, cellular PulG levels were reduced in the absence of PulL or PulM (Fig. 3B).

### 3.3. The periplasmic coiled coil is required for assembly of the PulL-PulM complex

To determine whether the deletion of periplasmic helices affects the PulL - PulM interaction, we used the bacterial two-hybrid (BACTH) assay (Karimova et al., 1998). Full-length PulL, PulM and their variants were fused to the C-termini of T18 and T25 CyaA fragments (Table 1). Constructs were co-transformed in *E. coli cya* mutant strain DHT1 to evaluate the reconstitution of adenylyl cyclase activity. By measuring the activity of beta-galactosidase activity, we compared interactions of full-length PulL and PulM constructs described previously (Dazzoni et al., 2022) and the PulL^ΔCC^ and PulM^ΔCC^ variants. For each tested couple, activity was measured for 8 independent cotransformants picked at random (Fig. 4).

**Figure 4.**
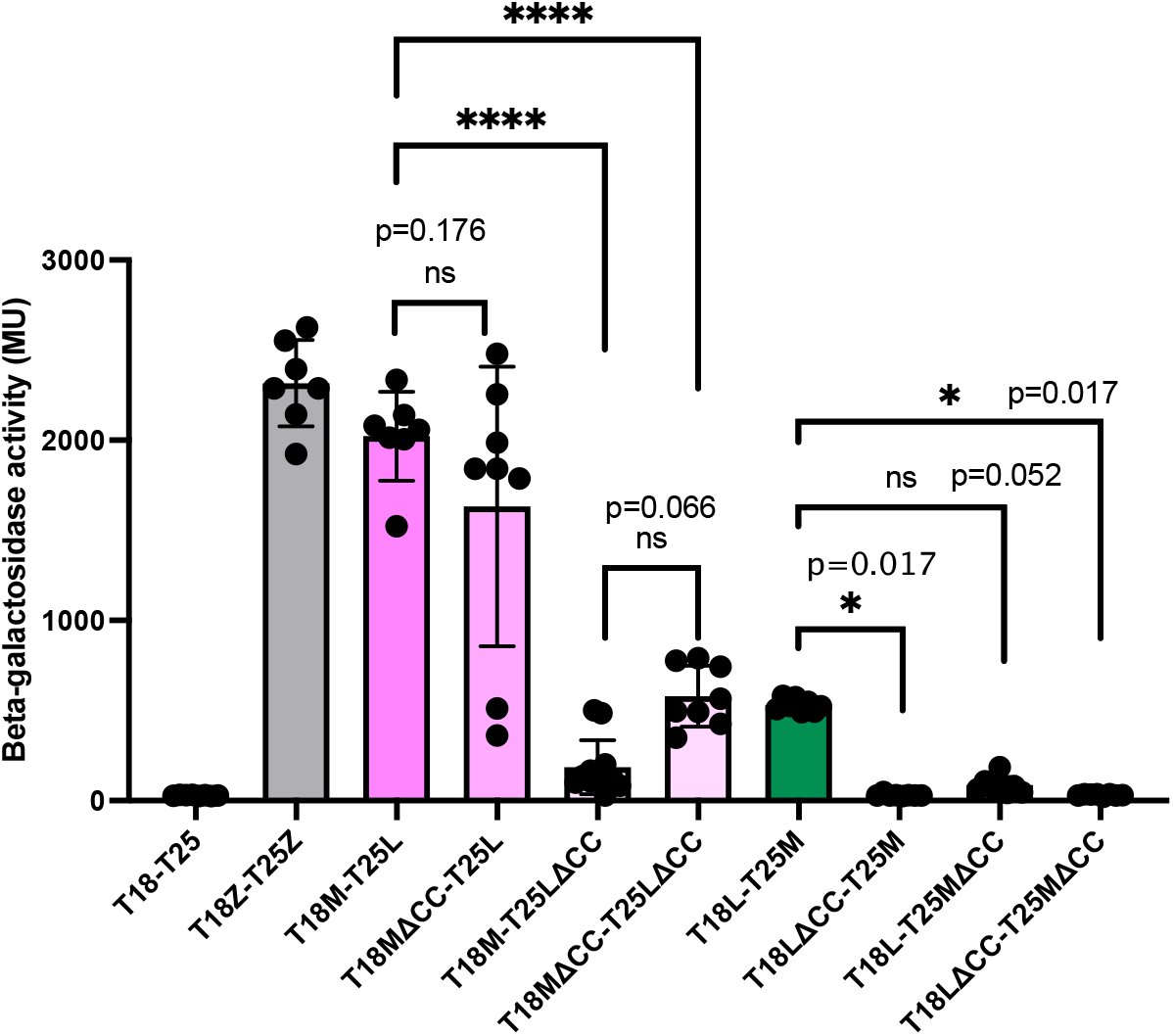
Periplasmic helices are required for PulL – PulM interaction. Beta-galactosidase activity (Miller units) of DHT1 bacteria co-transformed with plasmids producing T18 and T25 or their hybrids as indicated: Z, yeast leucin zipper; M (PulM), L (PulL); MΔCC (PulM^ΔCC^) and LΔCC (PulL^ΔCC^). The bar graph heights indicate mean values and error bars show standard deviation. Black dots show β-gal. activities of individual cultures. Statistical analysis was performed with GraphPad Prism 9 using One-way ANOVA test with multiple comparisons. ****, p<0.0001. Exact p values are indicated above the bars.

Bacteria containing plasmids encoding T18-PulM and T25-PulL showed high beta-galactosidase activity, comparable to the yeast leucin zipper positive control. In strains harboring T18-PulM^ΔCC^ and T25-PulL, the mean activity remained high, suggesting an interaction of PulM^ΔCC^ similar to that of PulM^WT^. However, in several isolates the activity was strongly reduced, indicating that removing the periplasmic helix affected PulM interaction with PulL. Deleting the PulL helix in T25-PulL^ΔCC^ had a much stronger effect and nearly abolished the interaction with T18-PulM. In comparison, combining the two deletions in T18-PulM^ΔCC^ and T25-PulL^ΔCC^ increased the mean activity, presumably by placing the respective CTDs at the same level relative to the membrane to facilitate their contacts. However, this increase was weak and not statistically significant in this experimental context.

Strains harboring T18-PulL and T25-PulM hybrids showed weaker β-galactosidase activity overall. This difference is probably due to the lower stability of T18-PulL compared to T18-PulM hybrids. Beta-galactosidase activity was reduced to background levels upon removal of the periplasmic helix residues in T18-PulL^ΔCC^ or T25-PulM^ΔCC^, and combining the two deletions in this context did not improve the interaction. Overall, these data suggest that coiled coil regions play a major role in the PulL-PulM assembly.

### 3.4. Periplasmic helix of PulM is not required for binding to PulG

PulM has been characterized as a targeting factor for PulH and PulG subunits during pseudopilus elongation (Nivaskumar et al., 2016, Santos-Moreno et al., 2017). Since the deletion of the PulM periplasmic helix affected pseudopilus assembly, we asked whether it also affects interactions with PulG. As shown in the BACTH assay results (Fig. 5), although deletion of the PulM periplasmic helix reduced the interaction of the pair T18-PulM – T25-PulG, the PulM^ΔCC^ variant could still interact with PulG strongly and significantly relative to the negative control. In addition, there was no effect on the interaction with T18-PulG.

**Figure 5.**
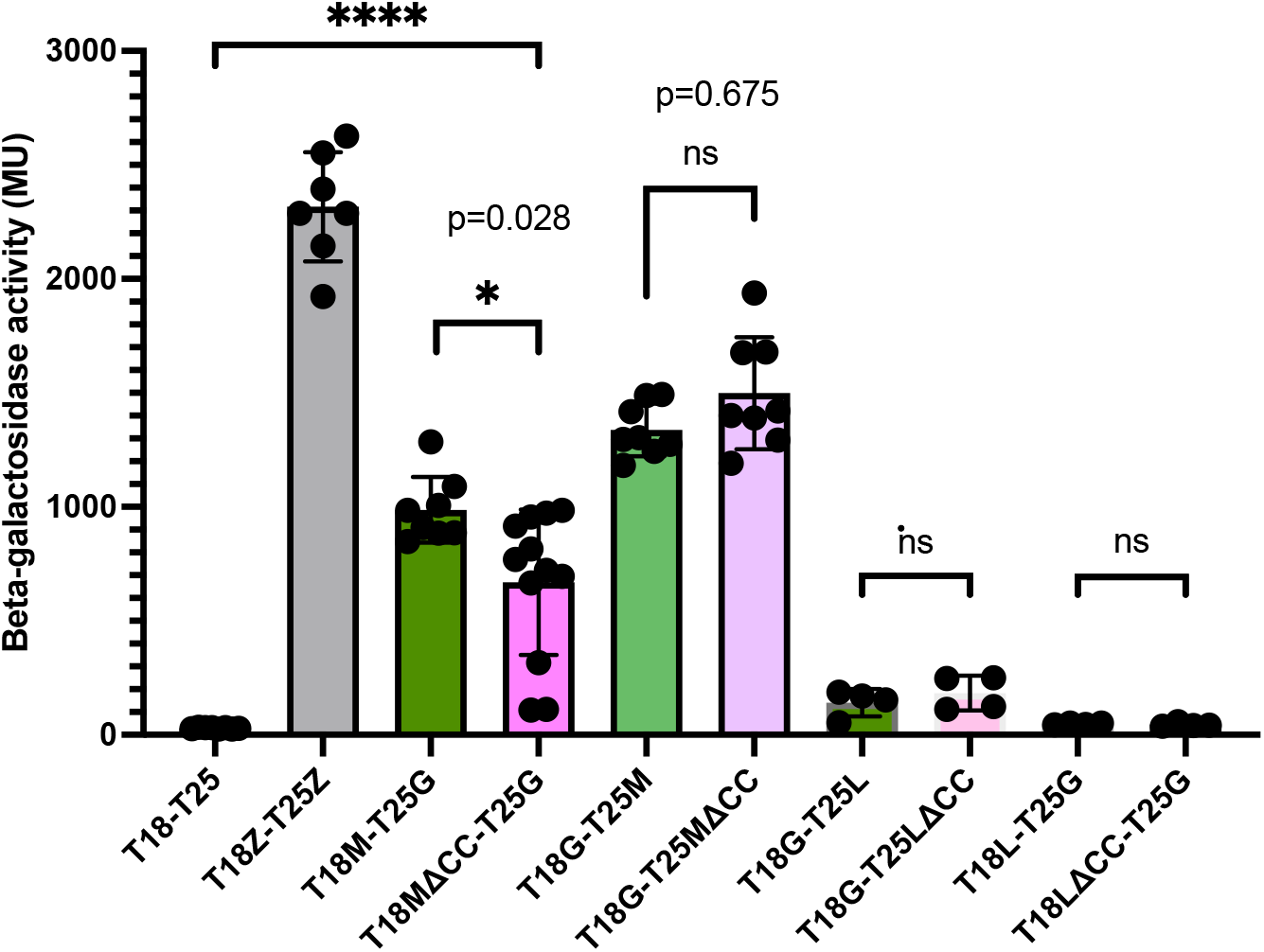
PulM periplasmic a-helix is not essential for interaction with PulG. Beta galactosidase activity measured for indicated pairs of T18- and T25-hybrids. Z, yeast leucin zipper; M (PulM), L (PulL); MΔCC (PulM^ΔCC^), LΔCC (PulL^ΔCC^) and G (PulG). Bar graphs indicate mean values and error bars standard deviation. Black dots show activity values of independent cultures. The data were plotted and analysed with GraphPad Prism 9 software.

Previous studies have shown that PulL does not interact with PulG in the BACTH assay (Nivaskumar et al., 2016). We confirmed this result here, by showing that β-galactosidase activities of the bacteria co-expressing *pulG* and *pulL* BACTH constructs were at the level of the negative control. Like the PulL^WT^, the PulL^ΔCC^ variant did not interact with PulG (Fig. 5). Based on these data, we concluded that it is unlikely that the piliation defect caused by PulM^ΔCC^ is due to a defect in pseudopilin binding.

### 3.5. The role of the cytoplasmic PulM peptide

The cytoplasmic domain of PulL interacts stably with the ATPase PulE, based on the study of their homologues in *Vibrio cholerae* (Abendroth et al., 2005). The cytoplasmic region of PulM is a 16-residue peptide, presumably functioning as a positively charged membrane anchor. We hypothesized that this region interacts with the polar N-terminal residues of PulG, as their substitutions (PulG^T2A^ and PulG^E5A^) strongly reduce interactions with PulM (Nivaskumar et al., 2016, Santos-Moreno et al., 2017). To test this hypothesis, we deleted the cytoplasmic peptide of PulM, and studied interactions of this variant designated PulMΔN with PulL and PulG. In the BACTH assay, PulMΔN variant showed a strong and significant defect in binding to PulG, confirming our predictions (Fig. 6A). This defect was comparable to that of PulM binding to the PulG^E5A^ variant, characterized previously (Santos-Moreno et al., 2017). At the same time, deleting the PulM cytoplasmic peptide did not affect its interactions with PulL in this assay.

**Figure 6.**
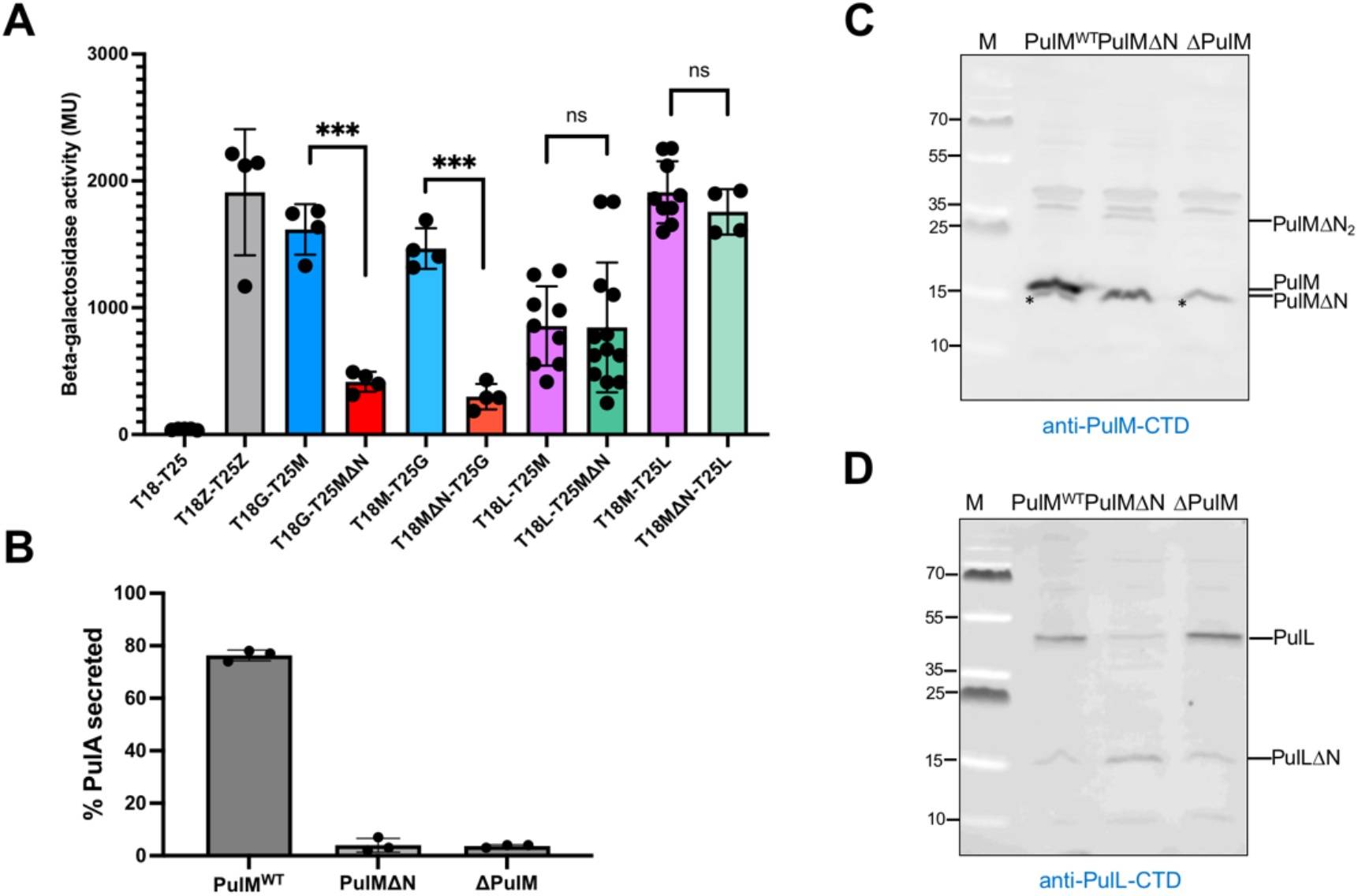
PulM cytoplasmic peptide is essential for interaction with PulG and for the PulL stability. **A**. BACTH analysis of PulM and PulMΔN interactions with PulG and PulL. **B**. PulA secretion in the presence of PulMΔN is fully defective. Each data point represents % of PulA secretion in strain PAP5378 containing plasmid pCHAP8496 (*ΔpulM*) complemented with *pulM*^WT^ (on pCHAP1353), *pulMAN* (on pCHAP8882) and empty vector (pSU18). Secretion assay was performed as described in Materials and Methods. (**C**) PulM (top) and (**D**) PulL (bottom) levels in strain PAP7460 carrying pCHAP8496 (*ΔpulM*) complemented with *pulM*^WT^ (on pCHAP1353), empty vector (pSU18) and *pulMΔN* (on pCHAP8882). The asterisk in (**C**) marks a nonspecific band cross-reacting with the antibody.

We tested the ability of PulMΔN to promote PulA secretion and found that nearly all PulA remained cell bound in the secretion assay, comparable to the negative control lacking PulM (Fig. 6B). This is probably caused by the defective interaction of PulMΔN with PulG. Although the levels of PulMΔN variant were reduced compared to PulM^WT^, the protein was still well produced and had the expected apparent size (~16.5 kDa) (Fig. 6C). Interestingly, this variant seemed to form some SDS-resistant dimers that were not detectable with PulM^WT^. We also tested the levels of PulL in these strains. Surprisingly, although the N-peptide deletion of PulM did not affect its interaction with PulL in the BACTH assay, we found that PulL was strongly destabilized in the presence of PulMΔN, resulting in a proteolytic product of a size similar to that of PulM (Fig. 6D). Antibodies directed against the PulL^CTD^ detected this product, suggesting that it comprises the inner membrane and periplasmic regions of PulL. As it appears to be generated by a specific cleavage that removes the PulL cytoplasmic domain, we designated this protein PulLΔN. These results suggest that the PulM cytoplasmic peptide not only ensures PulG targeting to the assembly site through direct interaction, but also protects PulL from proteolysis.

## 4. Discussion

Structural information on the PulL_CTD_-PulM_CTD_ heterodimer, together with BACTH and cysteine crosslinking data, allowed Dazzoni and collaborators to propose a model of membrane-embedded PulL-PulM complex (Fig. 1A) (Dazzoni et al., 2022). Sequence-based predictions (Ludwiczak, Winski et al., 2019) and biochemical data (Dazzoni et al., 2022) suggest that periplasmic helices of PulL and PulM interact together to form a coiled coil. Here we describe deletion, interaction and functional studies that support this model and show that this coiled coil plays a major role in PulL-PulM assembly. Deleting residues 270-302 of PulL abolished interaction with PulM in the BACTH assay, and removal of PulM residues 37-68 reduced or abolished PulL binding, depending on the plasmid construct and expression levels. In structural models of PulL and PulM, these deletions would shorten their periplasmic helices by 4.8 nm. This might preclude interaction of ferredoxin-like CTDs between the deletion variant and the native, full-length partner (Fig. 7A). Consistently, the BACTH interaction signal was somewhat improved when the two deletions were combined in T18-PulM^ΔCC^ and T25-PulL^ΔCC^. However, their interaction remained weak overall. The binding of PulL_CTD_ to PulM_CTD_ is weak and dynamic in solution (Dazzoni et al., 2022). Our results show that this interaction remains weak even when the two domains are anchored in the membrane *via* transmembrane segments. This suggests that the main role of the coiled coil is to stimulate PulM binding to PulL.

When isolated and in solution, PulL_CTD_ and PulM_CTD_ can form different types of parallel and anti-parallel homodimers, as shown by X-ray crystallography (Dazzoni et al., 2022). Similar behavior was found for the CTDs of EpsM, the PulM homologue from *V. cholerae* (Abendroth, Rice et al., 2004) and XcpL, the PulL homologue from *P. aeruginosa* (Fulara, Vandenberghe et al., 2018). Likewise, homo-oligomerization was observed for ferredoxin-like domains of T4P assembly proteins PilN and PilO, which are considered as orthologues of PulL transmembrane and periplasmic regions and of PulM, respectively. The CTD of PilN from *Thermus thermophilus* (Karuppiah, Collins et al., 2013) and soluble periplasmic regions of *P. aeruginosa* PilO and PilN formed homodimers in crystals (Sampaleanu, Bonanno et al., 2009). This is in contrast to their poor oligomerization *in vivo* in the bacterial membrane, which reduces the degrees of freedom for CTDs binding. Thus, the BACTH data indicate that PulL and PulM preferentially form heterodimers, with a parallel orientation, compatible with their membrane insertion (Dazzoni et al., 2022). Similarly, full-length *P. aeruginosa* PilN and PilO preferentially formed heterodimers, and no homodimers were observed in the BACTH assay (Leighton, Dayalani et al., 2015). The long periplasmic helices thus seem to provide an interacting surface which guides this coiled coil formation making it more efficient and accurate.

In the *Dickeya dadanti* T2SS, the periplasmic regions of OutL and OutM also comprise the major interacting interfaces as shown in BACTH and copurification studies (Lallemand et al., 2013). However, unlike in the Pul system, the ferredoxin-like domains of OutL and OutM appear to interact more strongly with each other than their larger periplasmic regions, suggesting a minor role for the coiled coil. Interestingly, mutations in the coiled coils of PilN and PilO, components of the *P. aeruginosa* T4P assembly system, caused defective pilus retraction (Leighton, Yong et al., 2016). Both studies concluded that precise and dynamic interactions between assembly platform components control the activity of the respective systems.

Deletions of the coiled coil regions strongly affected PulL and PulM functions. Although variant PulM^ΔCC^ was more stable and retained the capacity to interact with PulL in the BACTH assay, it was as defective as PulL^ΔCC^ for both PulA secretion and PulG pilus assembly. This may be due to impaired structure and function of the PulL-PulM complex. Coiled coils might provide structural rigidity to the complex compared to the individual proteins. Shortening of PulL and PulM helices by 4.8 nm could also result in defective interactions with other T2SS components or cellular structures, such as the peptidoglycan layer. In PulL^ΔCC^ and PulM^ΔCC^, the globular ferredoxin-like domains (CTDs) would be placed proximal to the membrane, connected *via* the flexible linkers to the transmembrane helices. This might affect the overall architecture of the T2SS complex and interactions with its substrates.

While we showed here that PulM^ΔCC^ retained its ability to bind PulG, the next step in pseudopilus assembly, possibly involving binding to PulL, might be impaired. Like other pilins and pseudopilins, the TM segment of PulG is alpha-helical in its membrane-embedded state, but undergoes stretching upon incorporation into the pseudopilus (López-Castilla et al., 2017). This results in the loss of secondary structure in the region between Gly14 and Pro22 of PulG, which adopts an extended coil conformation (López-Castilla et al., 2017). The force that causes this stretching is likely to be generated during PulG extraction from the membrane, a key step in pseudopilus assembly which involves binding to PulM (Santos-Moreno et al., 2017). The PulL-PulM coiled coil might act as an anchoring point during this step. Rigidity or the length of PulL-PulM dimer might be essential to exert force on PulG and operate its stretching during membrane extraction. The PulE ATPase in complex with the PulL cytoplasmic domain might be directly involved in this step. A defect in membrane extraction step could cause the major piliation defect of *pulL^ΔCC^* and *pulM^ΔCC^* mutants observed here.

Our BACTH data suggest that the PulM cytoplasmic region in not required for interaction with PulL, and that the periplasmic regions play a dominant role. The specific cleavage of PulL cytoplasmic domain in the *pulMΔN* mutant was therefore surprising. We hypothesize that the N-terminal peptide of PulM protects PulL from proteolysis by binding to a disordered linker that connects PulLcyto domain with the fragment comprising PulL transmembrane and periplasmic regions. This linker is present in the PulL model from the AF2 database (Varadi, Anyango et al., 2022) predicted by the Alpha Fold 2 algorithm (Jumper, Evans et al., 2021) (Fig. S1).

Binding to the disordered PulL linker might require the free N-terminus of PulM, and thus might occur in the native context, but not in the BACTH constructs, where PulM N-terminus is fused to the T18 and T25 CyaA fragments. A similar interaction is observed in T4P assembly systems, between the conserved N-terminal peptide of PilN and PilM which is a paralogue of the PulL_cyto_ domain (Karuppiah & Derrick, 2011) (McCallum, Tammam et al., 2016). This contact is highly important for pilus assembly (Georgiadou, Castagnini et al., 2012) and involves a conserved sequence ΦNLLP (where Φ is a V, I or L) at the N-terminus of PilN homologues. Structural studies have shown that this N-terminal peptide fits into a groove in PilM to form a stable PilM-PilN complex – a functional equivalent of PulL. The conserved Asn residue (N5 in PilN of *T. thermophilus*) forms a network of hydrogen bonds with several residues in the PilM groove, suggesting a major role in PilM binding (Karuppiah & Derrick, 2011). Functional data in *Neisseria meningitidis* support the crucial role of Asn8 in PilN showing a strong piliation defect of the PiIN_N8A_ variant (Georgiadou et al., 2012). The presence of a similar N-terminal sequence in PulM, MH**NLL** (identical residues shown in bold) led us to test the effect of N3A substitution in PulM. The PulM^N3A^ variant showed a strong secretion defect and affected the stability of both PulM and PulL (Dazzoni et al., 2022). These effects were reproduced in the PulMΔN variant described here. The *pulMΔN* mutants show reduced PulM and PulL stability, generating a distinct PulL fragment corresponding in size and probably in domain organization to the PilN component of T4P assembly systems. Of note, a similar PulL degradation product is also observed in the presence of native PulM (Dazzoni et al., 2022), suggesting that the cytoplasmic linker region of PulL is dynamic and protease sensitive, even in the presence of its interacting partner(s).

The above results argue that, although PulM is often considered as a paralogue of PilO in T4P systems, its role may be more similar to that of PilN. Importantly, direct interaction of PulM with the major pseudopilin PulG parallels the binding of the PilN homologue HofN to the major pilin PpdD in the T4P assembly system of Enterohaemorrhagic *E. coli* (Luna Rico, Zheng et al., 2019).

The N-terminal regions of PulM and PilN might therefore play analogous roles in pilin targeting to the assembly site. The phenotype of the PulMΔN variant, strongly defective in PulG binding and affecting PulL stability, suggests a role of the cytoplasmic PulM peptide in passing the pseudopilin subunits to PulL for incorporation into the growing pilus. PulM is also more abundant in the cell relative to PulL (Dazzoni et al., 2022) consistent with its role as a shuttle between the large membrane pool of free PulG and PulL stably bound to the ATPase PulE at the pseudopilus assembly site. Crosslinking studies of *V. cholerae* T2SS provided evidence for a transient complex of EpsL with EpsG (Gray, Bagdasarian et al., 2011), supporting the formation of G-M-L tripartite complex. Clearly, further studies are needed to identify their precise interactions within this complex and conformational changes of partners that promote this key step during pseudopilus assembly.

## Acknowledgments

We thank N. Izadi-Pruneyre, I. Guilvout, E. Bouveret and members of the BIM unit for helpful discussions. YL was funded by the Pasteur Paris University (PPU) international PhD program and the China National Biotec Group Company Limited, and by a doctoral fellowship from the China Scholarship Council. JSM was funded by a fellowship from the Basque Government. This work was funded by Institut Pasteur, CNRS and ANR grant Synergy-T2SS ANR-19-CE11-0020-01.

## References

Abendroth J, Murphy P, Sandkvist M, Bagdasarian M, Hol WG (2005) The X-ray structure of the type II secretion system complex formed by the N-terminal domain of EpsE and the cytoplasmic domain of EpsL of *Vibrio cholerae*. J Mol Biol 348: 845–55.

Abendroth J, Rice AE, McLuskey K, Bagdasarian M, Hol WG (2004) The crystal structure of the periplasmic domain of the type II secretion system protein EpsM from *Vibrio cholerae:* the simplest version of the ferredoxin fold. J Mol Biol 338: 585–96.

Campos M, Nilges M, Cisneros M, Francetic O (2010) Detailed structure and assembly model of the type II secretion pilus from sparse data. Proc Natl Acad Sci USA 107: 13081–6.

Chernyatina AA, Low HH (2019) Core architecture of a bacterial type II secretion system. Nat Commun 10: 5437.

Cianciotto NP, White RC (2017) Expanding Role of Type II Secretion in Bacterial Pathogenesis and Beyond. Infect Immun 85: e00014-17

Cisneros DA, Bond PJ, Pugsley AP, Campos M, Francetic O (2012) Minor pseudopilin self-assembly primes type II secretion pseudopilus elongation. The EMBO J 31: 1041–1053.

Datsenko KA, Wanner BL (2000) One-step inactivation of chromosomal genes in *Escherichia coli* K-12 using PCR products. Proc Natl Acad Sci U S A 97: 6640–6645.

Dautin N, Karimova G, Ullmann A, Ladant D (2000) Sensitive Genetic Screen for Protease Activity Based on a Cyclic AMP Signaling Cascade in *Escherichia coli*. J Bacteriol 182: 7060–7066.

Dazzoni R, Li Y, Lopez-Castilla A, Brier S, Mechaly A, Cordier F, Haouz A, Nilges M, Francetic O, Bardiaux B, Izadi-Pruneyre N (2022) Structure and dynamic association of an assembly platform subcomplex of the bacterial type II secretion system. BioRXiv https://doi.org/10.1101/2022.07.16.500195.

Douzi B, Ball G, Cambillau C, Tegoni M, Voulhoux R (2011) Deciphering the Xcp *Pseudomonas aeruginosa* Type II Secretion Machinery through Multiple Interactions with Substrates. J Biol Chem 286: 40792–40801.

Durand E, Bernadac A, Ball G, Lazdunski A, Sturgis JN, Filloux A (2003) Type II protein secretion in *Pseudomonas aeruginosa*: the pseudopilus is a multifibrillar and adhesive structure. J Bacteriol 185: 2749–58.

Escobar CA, Douzi B, Ball G, Barbat B, Alphonse S, Quinton L, Voulhoux R, Forest KT (2021) Structural interactions define assembly adapter function of a type II secretion system pseudopilin. Structure 29: 1116–1127.e8.

Fulara A, Vandenberghe I, Read RJ, Devreese B, Savvides SN (2018) Structure and oligomerization of the periplasmic domain of GspL from the type II secretion system of *Pseudomonas aeruginosa*. Sci Rep 8: 16760

Georgiadou M, Castagnini M, Karimova G, Ladant D, Pelicic V (2012) Large-scale study of the interactions between proteins involved in type IV pilus biology in *Neisseria meningitidis:*characterization of a subcomplex involved in pilus assembly. Mol Microbiol 84: 857–873.

Gray MD, Bagdasarian M, Hol WGJ, Sandkvist M (2011) In vivo cross-linking of EpsG to EpsL suggests a role for EpsL as an ATPase-pseudopilin coupling protein in the Type II secretion system of *Vibrio cholerae*. Mol Microbiol 79: 786–798.

Gu S, Shevchik VE, Shaw R, Pickersgill RW, Garnett JA (2017) The role of intrinsic disorder and dynamics in the assembly and function of the type II secretion system. Biochim Biophys Acta (BBA) - Proteins and Proteomics 1865: 1255–1266.

Hay ID, Belousoff MJ, Dunstan RA, Bamert RS, Lithgow T (2018) Structure and membrane topography of the vibrio-type secretin complex from the type 2 secretion system of enteropathogenic *Escherichia coli*. J Bacteriol 200: e00521–17.

Jumper J, Evans R, Pritzel A, Green T, Figurnov M, Ronneberger O, Tunyasuvunakool K, Bates R, Žídek A, Potapenko A, Bridgland A, Meyer C, Kohl SAA, Ballard AJ, Cowie A, Romera-Paredes B, Nikolov S, Jain R, Adler J, Back T et al. (2021) Highly accurate protein structure prediction with AlphaFold. Nature 596: 583–589.

Karimova G, Pidoux J, Ullmann A, Ladant D (1998) A bacterial two-hybrid system based on a reconstituted signal transduction pathway. Proc Natl Acad Sci U S A 95: 5752–5756.

Karuppiah V, Collins RF, Thistlethwaite A, Gao Y, Derrick JP (2013) Structure and assembly of an inner membrane platform for initiation of type IV pilus biogenesis. Proc Nati Acad Sci USA 110: E4638–E4647.

Karuppiah V, Derrick JP (2011) Structure of the PilM-PilN Inner Membrane Type IV Pilus Biogenesis Complex from *Thermus thermophilus*. J Biol Chem 286: 24434–24442.

Kohler R, Schafer K, Muller S, Vignon G, Diederichs K, Philippsen A, Ringler P, Pugsley AP, Engel A, Welte W (2004) Structure and assembly of the pseudopilin PulG. Mol Microbiol 54: 647–64.

Korotkov K, Sandkvist M (2019) Architecture, Function, and Substrates of the Type II Secretion System. EcoSal Plus 8(2) 10.1128/ecosalplus.ESP-0034-2018.

Korotkov KV, Hol WG (2008) Structure of the GspK-GspI-GspJ complex from the enterotoxigenic *Escherichia coli* type 2 secretion system. Nature structural & molecular biology 15: 462–8.

Lallemand M, Login FH, Guschinskaya N, Pineau C, Effantin G, Robert X, Shevchik VE (2013) Dynamic Interplay between the Periplasmic and Transmembrane Domains of GspL and GspM in the Type II Secretion System. PLoS One 8: e79562

Leighton TL, Dayalani N, L.M. S, Howell PL, LL. B (2015) Novel Role for PilNO in Type IV Pilus Retraction Revealed by Alignment Subcomplex Mutations. J Bacteriol 197: 2229–2238.

Leighton TL, Yong DH, Howell PL, Burrows LL (2016) Type IV Pilus Alignment Subcomplex Proteins PilN and PilO Form Homo- and Heterodimers *n Vivo*. J Biol Chem 291: 19923–19938.

López-Castilla A, Thomassin J-L, Bardiaux B, Zheng W, Nivaskumar M, Yu X, Nilges M, Egelman EH, Izadi-Pruneyre N, Francetic O (2017) Structure of the calcium-dependent type 2 secretion pseudopilus. Nat Microbiol 2: 1686–1695.

Ludwiczak J, Winski A, Szczepaniak K, Alva V, Dunin-Horkawicz S (2019) DeepCoil—a fast and accurate prediction of coiled-coil domains in protein sequences. Bioinformatics (Oxford, England) 35: 2790–2795.

Luna Rico A, Zheng W, Petiot N, Egelman EH, Francetic O (2019) Functional reconstitution of the type IVa pilus assembly system from enterohaemorrhagic *Escherichia coli*. Mol Microbiol 111: 732–749.

McCallum M, Tammam S, Little DJ, Robinson H, Koo J, Shah M, Calmettes C, Moraes TF, Burrows LL, Howell PL (2016) PilN Binding Modulates the Structure and Binding Partners of the *Pseudomonas aeruginosa* Type IVa Pilus Protein PilM*. J Biol Chem 291: 11003–11015.

Michaelis S, Chapon C, d’Enfert C, Pugsley AP, Schwartz M (1985) Characterization and expression of the structural gene for pullulanase, a maltose-inducible secreted protein of *Klebsiella pneumoniae*. J Bacteriol 164: 633–638.

Miller JH (1972) Experiments in Molecular Genetics. Cold Spring Harbor Laboratory, Cold Spring Harbor, NY. USA,

Naskar S, Hohl M, Tassinari M, Low HH (2021) The structure and mechanism of the bacterial type II secretion system. Mol Microbiol 115: 412–424.

Nivaskumar M, Bouvier G, Campos M, Nadeau N, Yu X, Egelman EH, Nilges M, Francetic O (2014) Distinct Docking and Stabilization Steps of the Pseudopilus Conformational Transition Path Suggest Rotational Assembly of Type IV Pilus-like Fibers. Structure 22: 685-696

Nivaskumar M, Santos-Moreno J, Malosse C, Nadeau N, Chamot-Rooke J, Tran Van Nhieu G, Francetic O (2016) Pseudopilin residue E5 is essential for recruitment by the type 2 secretion system assembly platform. Mol Microbiol 101: 924–941.

Patrick M, Korotkov KV, Hol WGJ, Sandkvist M (2011) Oligomerization of EpsE Coordinates Residues from Multiple Subunits to Facilitate ATPase Activity. J Biol Chem 286: 10378–10386.

Possot O, Vignon G, Bomchil N, Ebel F, Pugsley AP (2000) Multiple interactions between pullulanase secreton components involved in stabilization and cytoplasmic membrane association of PulE. J Bacteriol 182: 2142–2152.

Py B, Loiseau F, Barras F (2001) An inner membrane platform in the type II secretion machinery of Gram-negative bacteria. EMBO Rep 2: 244–248.

Robien MA, Krumm BE, Sandkvist M, Hol WG (2003) Crystal structure of the extracellular protein secretion NTPase EpsE of *Vibrio cholerae*. J Mol Biol 333: 657–74.

Sampaleanu LM, Bonanno JB, Ayers M, Koo J, Tammam S, Burley SK, Almo SC, Burrows LL, Howell PL (2009) Periplasmic Domains of *Pseudomonas aeruginosa* PilN and PilO Form a Stable Heterodimeric Complex. J Mol Biol 394: 143–159.

Sandkvist M, Hough LP, Bagdasarian MM, Bagdasarian M (1999) Direct interaction of the EpsL and EpsM proteins of the general secretion apparatus in *Vibrio cholerae*. J Bacteriol 181: 3129–35.

Santos-Moreno J, East A, Guilvout I, Nadeau N, Bond PJ, Tran Van Nhieu G, Francetic O (2017) Polar N-terminal Residues Conserved in Type 2 Secretion Pseudopilins Determine Subunit Targeting and Membrane Extraction Steps during Fibre Assembly. J Mol Biol 429: 1746–1765.

Sauvonnet N, Gounon P, Pugsley AP (2000) PpdD type IV pilin of *Escherichia coli* K-12 can be assembled into pili in *Pseudomonas aeruginosa*. J Bacteriol 182: 848–854.

Sauvonnet N, Vignon G, Pugsley AP, Gounon P (2000) Pilus formation and protein secretion by the same machinery in *Escherichia coli*. EMBO J 19: 2221–2228.

Schägger H, von Jagow G (1987) Tricine-sodium dodecyl sulfate-polyacrylamide gel electrophoresis for the separation of proteins in the range from 1 to 100 kDa. Anal Biochem 166: 368–379.

Varadi M, Anyango S, Deshpande M, Nair S, Natassia C, Yordanova G, Yuan D, Stroe O, Wood G, Laydon A, Žídek A, Green T, Tunyasuvunakool K, Petersen S, Jumper J, Clancy E, Green R, Vora A, Lutfi M, Figurnov M et al. (2022) AlphaFold Protein Structure Database: massively expanding the structural coverage of protein-sequence space with high-accuracy models. Nucl Acids Res 50: D439–D444.

Yan Z, Yin M, Xu D, Zhu Y, Li X (2017) Structural insights into the secretin translocation channel in the type II secretion system. Nat Struct Mol Biol 24: 177–183.

