## Supplemental data for "Periplasmic coiled coil formed by assembly platform proteins PulL and PulM is critical for function of the *Klebsiella* type II secretion system"

**Supplementary Table S1**

**Supplementary Fig. 1**

**Table S1 Oligonucleotides used in this study**

| Primer name | DNA sequence (5'-3') |
|---|---|
| PulL dCC For | CCTCCACGATCATCTCATGCAGCAGCATCTGCGGCAG |
| PulL dCC Rev | GCTTCTGATTGCGAAGCTGCCATAACGTGTAATAG |
| PulM dCC For | CTATTACACGTTATGGCAGCTTCGCAATCAGAAGC |
| PulM dCC Rev | GCTTCTGATTGCGAAGCTGCCATAACGTGTAATAG |
| PulM del N5 | CTGACGCTGGAGGGGAACGATGCGGTGTCTGCTGCTGGGTATG |
| PulM del N3 | CATACCCAGCAGCAGACACCGCATCGTTCCCCTCCAGCGTCAG |
| PulM delN For | CGGTGTCTGCTGCTGGGTATG |
| PulM delN Rev2 | ATGCATCGGATCCTCTAGAGTC |

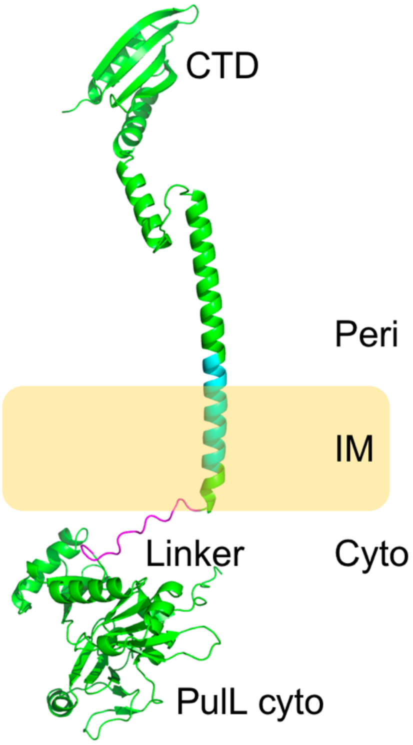


**Figure S1.** The AlphaFold 2 model of PulL in green represented in the bacterial inner membrane (IM), shown in yellow. The PulL cytoplasmic ($PulL_{cyto}$) and C-terminal domains (CTD) are labeled. The disordered linker is shown in magenta. The periplasmic (peri) and cytoplasmic (cyto) compartments are indicated.
